# Regional heterogeneity of developing GABAergic interneuron excitation *in vivo*

**DOI:** 10.1101/701862

**Authors:** Yasunobu Murata, Matthew T. Colonnese

## Abstract

GABAergic interneurons are proposed to be critical for early activity and synapse formation by directly exciting, rather than inhibiting, neurons in developing hippocampus and neocortex. However, the role of GABAergic neurons in the generation of neonatal network activity has not been tested *in vivo*, and recent studies have challenged the excitatory nature of early GABA. By locally manipulating interneuron activity in unanesthetized neonatal mice, we show that GABAergic neurons are indeed excitatory in hippocampus at postnatal-day 3 (P3), and responsible for most of the spontaneous firing of pyramidal cells at that age. Hippocampal interneurons become inhibitory by P7, whereas cortical interneurons are inhibitory at P3 and remain so throughout development. This regional and age heterogeneity is the result of a change in chloride reversal potential as activation of light-gated anion channels expressed in glutamatergic neurons causes firing in hippocampus at P3, but silences it at P7. This study in the intact brain reveals a critical role for GABAergic interneuron excitation in neonatal hippocampus, and a surprising heterogeneity of interneuron function in cortical circuits that was not predicted from *in vitro* studies.

## Main Text

The developing brain spontaneously generates region- and age-specific activity, which is fundamental to circuit formation (Kirkby et al., 2013). Many of these early activities are present at ages when low expression of the neuronal K-Cl co-transporter KCC2 renders GABAA receptors depolarizing, and sometimes excitatory, in excised tissue preparations (Rivera et al., 1999). This early excitatory action of GABAergic neurons, primarily mediated by medial ganglion eminence derived somatostatin-expressing neurons in hippocampus drives synchronized activity *in vitro* (Flossmann et al., 2019; Wester and McBain, 2016), and supports glutamatergic and GABAergic synapse formation in cortical circuits (Ben-Ari et al., 2007; Oh et al., 2016). Excitatory GABA has further been hypothesized to be critical to generation of early synchronized activity *in vivo* in human neonates and animals, with the switch to inhibitory GABA causing loss of these early patterns (Le Magueresse and Monyer, 2013; Myers et al., 2012; Vanhatalo and Kaila, 2006). Understanding the nature of GABAergic signaling is essential to developing treatments for seizures in infants, in whom standard augmentation of GABAA receptor signaling is often ineffective (Glykys et al., 2017; Khazipov et al., 2015).

Despite the central importance of GABAergic circuits to brain development and function, direct tests *in vivo* of GABA’s excitatory nature and of the role of GABAergic neurons in early activity have been rare (Kirmse and Holthoff, 2017). Pharmacological blockade of GABA_A_ receptors in sensory cortex causes network bursting, suggesting an inhibitory action *in vivo* (Minlebaev et al., 2007, 2011), but such blockade has paradoxical effects in hippocampal slices in which GABA is clearly excitatory, making these studies non-definitive (Khalilov et al., 1999; Lamsa et al., 2000). Two recent studies were unable to demonstrate excitatory GABA *in vivo* even when it is excitatory at the same ages *in vitro* (Kirmse et al., 2015; Valeeva et al., 2016). In addition to the question of GABAergic polarity, the role of local interneurons themselves in the generation of early synchronized activity *in vivo* is unknown.

To directly test the role of local GABAergic neurons in cortical circuits--where seizures are most commonly generated and GABA is considered to remain excitatory late into development (Kirmse and Holthoff 2017)--for the generation of early spontaneous activity, we employed a variety of chemogenetic and optogenetic approach to manipulate the activity of GABAergic interneurons while recording network activity locally using multi-electrode array recordings in un-anesthetized neonatal mice. We first used local expression of the cre-dependent adeno-associated virus encoding the inhibitory kappa-opioid receptor Designer Receptors Exclusively Activated by Designer Drugs (KOR-DREADD or KORD) (Vardy et al., 2015) or the excitatory DREADD (hM3Dq) (Armbruster et al., 2007) in hippocampus or visual cortex of GAD2-cre mice (Taniguchi et al., 2011) to probe the role of interneuron activity. After local viral injection at P0, both KORD and hM3Dq could be detected by P3. In both structures, expression was limited to GABAergic neurons (Fig.1A-B & Fig.3A-B). Whole-cell current-clamp recordings in slices showed hyperpolarization by SalB (the KORD agonist) and depolarization by CNO (the hM3Dq agonist) in GABAergic neurons by P3, which was similar to P11 (Fig.1C & Fig.3C). Thus, this approach allowed for suppression (KORD-SalB) and enhancement (hM3Dq-CNO) of GABAergic neuronal activity in neonatal hippocampus and cortex.

**Fig. 1.**
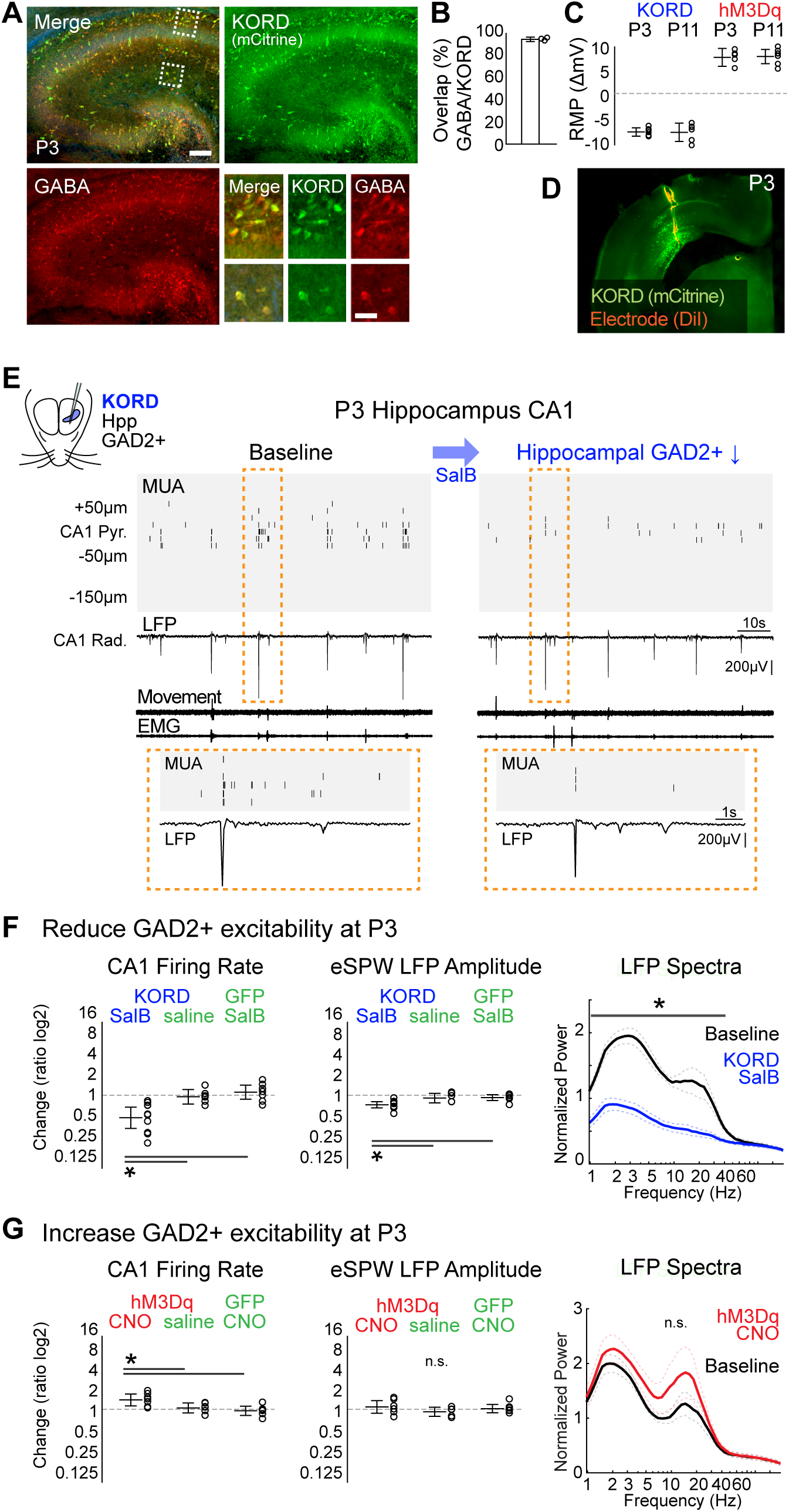
GABAergic interneurons are excitatory in 3-day old hippocampus *in vivo*. **(A)** Co-labeling of AAV-dF-KORD-IRES-mCitrine expressed in GAD2-Cre mouse with anti-GABA to label GABAergic neurons. Scale bars: 100μm and 20μm. **(B)** Percentage of KORD expressing neurons that also express GABA (n=3, mean+/− 95%CI, 94.7%±3.4%). **(C)** Change in membrane potential upon agonist perfusion for KORD or hM3Dq expressing CA1 neurons in slices at P3 and P11 (n=6,6,5,7; mean +/− 95%CI, −7.43±0.86, −7.5±1.89, 7.92±1.84, 8.09±1.46). **(D)** Representative localization of electrode and viral expression in P3 animal. **(E)** Representative recording for P3 reduction of GABAergic neuron excitability. Multi-unit spike activity (MUA) of spontaneous activity in CA1 hippocampus, along with associated *stratum radiatum* local field potential (LFP) and thoracic movement detection and electromyography (EMG). Activity is dominated by early sharp waves (eSPW) whose spike density is reduced following subcutaneous SalB (KORD agonist) injection. **(F)** Quantification of KORD-induced suppression of GABAergic neuron excitability and control conditions. From left: change in pyramidal cell layer firing rate (n=10,6,8; log2 ratio, mean +/− 95%CI: −1.23±0.58, −0.08±0.4, 0.17±0.39; ANOVA for manipulation, p<0.001), change in eSPW amplitude (−0.53±0.17, −0.16±0.26, −0.13±0.16; p=0.002), and normalized (to mean of 1-100Hz baseline) spectral power for *stratum radiatum* LFP (n=10, permutation test, p>0.05 at 1.0-48.7Hz). **(G)** Quantification of hM3Dq induced increase in GABAergic excitability (n=7,6,6; 0.51±0.33, 0.07±0.28, −0.09±0.25; p=0.005; 0.13±0.35, −0.14±0.25, 0.03±0.23; p=0.33; n=7, n.s. for all Hz).

**Fig. 2.**
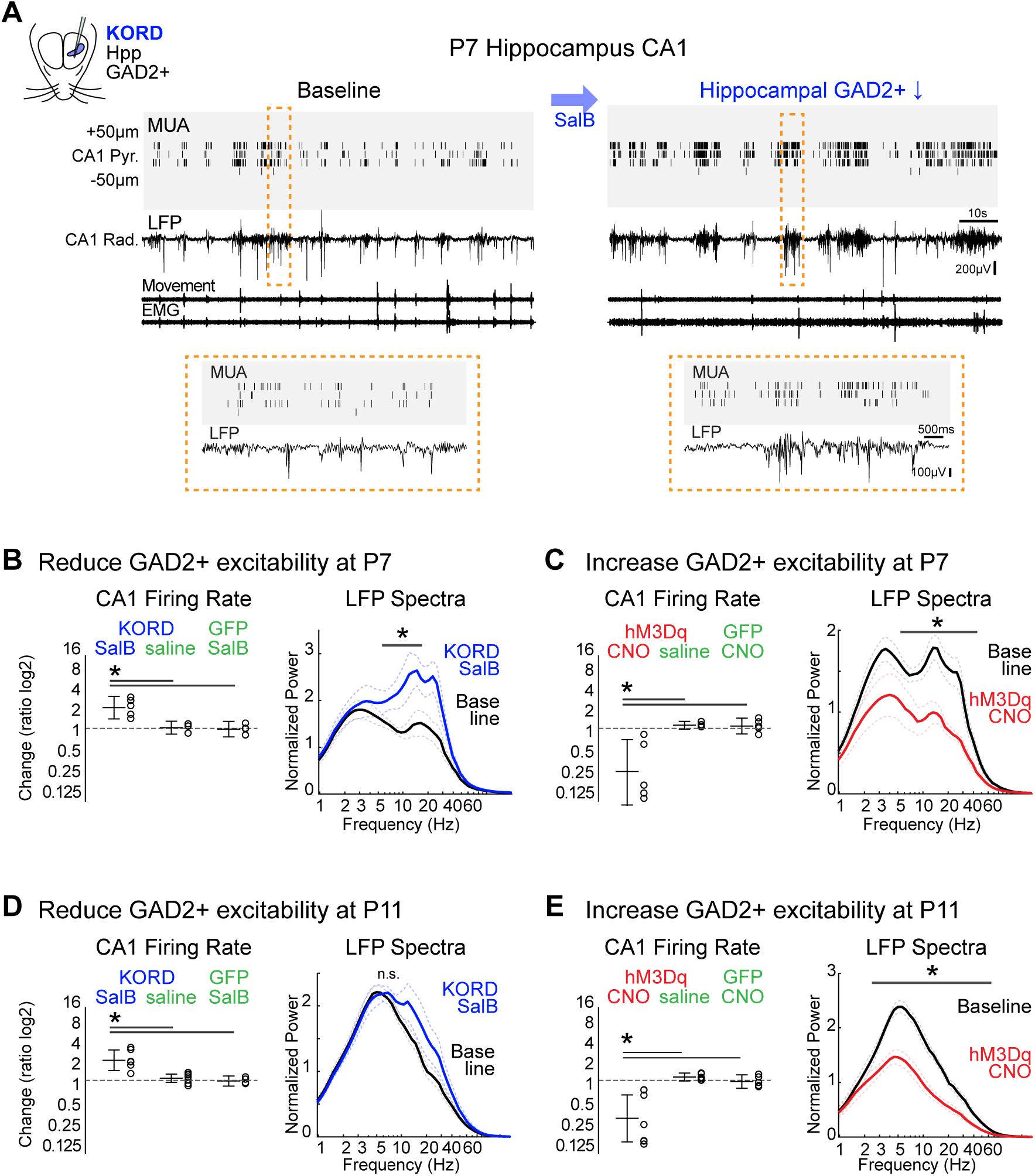
Hippocampal GABAergic neurons are inhibitory by P7. **(A)** Representative recording for GABAergic neuron suppression in P7 hippocampus. **(B-C)** Quantification of suppression (B) and enhancement (C) of GABAergic neuron excitability at P7 (B: n=5,4,4; 1.14±0.62, 0.04±0.35, −0.04±0.43; p=0.001; n=5, p<0.05 at 6.9-14.7 Hz; C: n=5,4,5; −2.37±2.02, 0.17±0.21, 0.13±0.43; p=0.003; n=5, p<0.05 at 7.7-93.3 Hz). **(D-E)** Similar quantification at P11 (D: n=6,8,4; 1.11±0.57, 0.12±0.22, −0.03±0.28; p>0.001; n=6, n.s; E: n=7,5,5; −2.09±1.29, 0.18±0.22, −0.07±0.37; p=0.001; n=7 p>0.05 at 2.3-93.3 Hz).

**Fig. 3.**
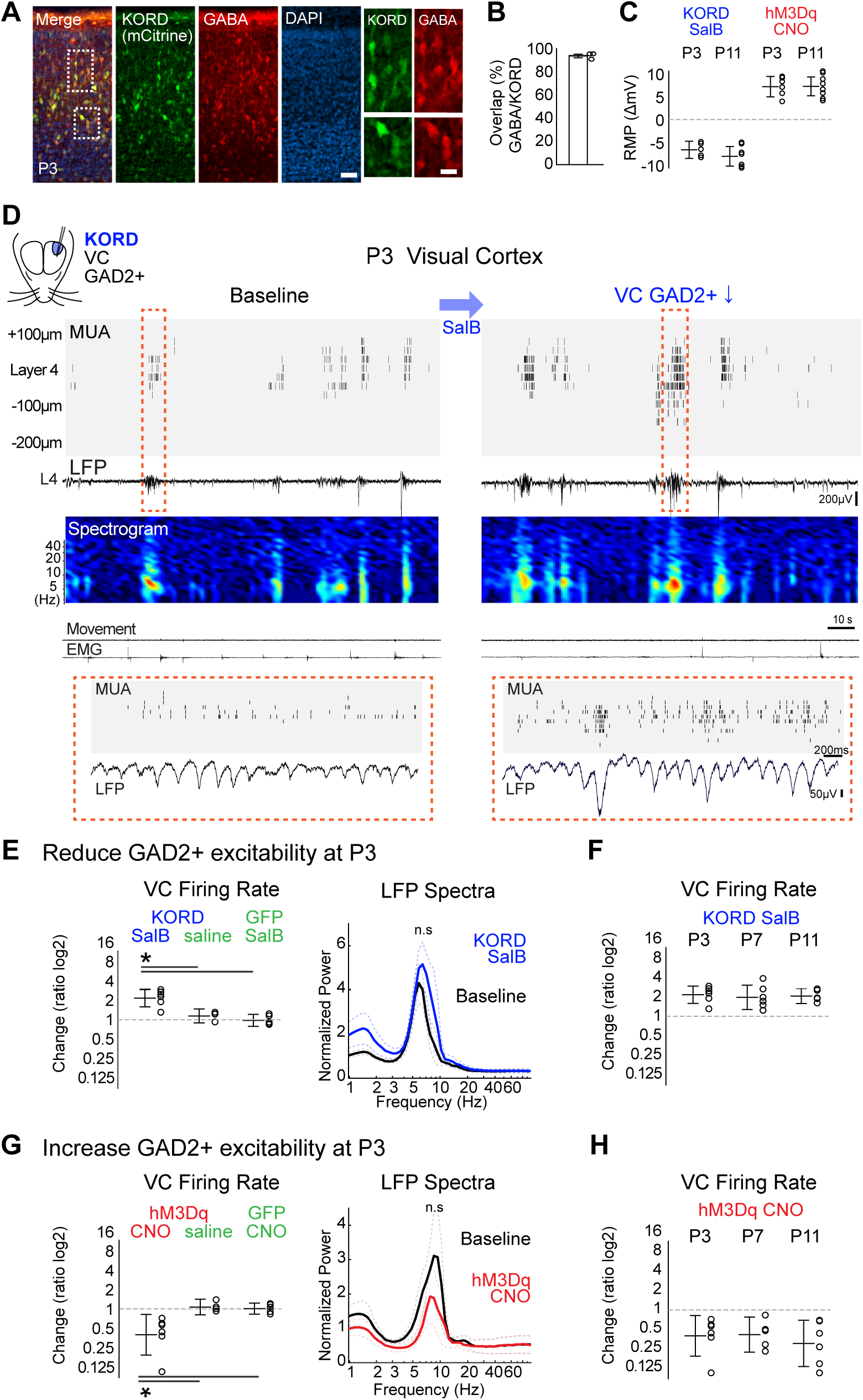
GABAergic neurons in visual cortex have a net inhibitory action as early as P3. **(A-B)** Co-localization of KORD expression with GABA (n=3, 93.7%±6.7%). Scale bars: 50μm and 10μm. **(C)** Change in membrane potential of GAD2+ neurons by activation of KORD and hM3Dq at P3 and P11 in visual cortical slices (n=5,7,6,8; - 6.62±1.88, −8.03±2.14, 7.18±2.23, 7.23±2.06). **(D)** Representative recording of visual cortex at P3 and the effect of GABAergic neuron suppression. LFP spectrogram is from presumptive layer 4. **(E)** Quantification of change in superficial layer firing rate (n=6,4,5; 1.25±0.5, 0.22±0.4, −0.03±0.35; p=0.001) and LFP spectral power (n=6; n.s.) following suppression of GABAergic neuron excitability by KORD activation. **(F)** Firing rate change at P3, P7 and P11 to KORD activation (n=6,6,5; 1.25±0.5, 1.11±0.71, 1.17±0.43; p=0.89). **(G-H)** As E-F but for GABAergic neuron enhancement by hM3Dq activation (G: n=6,4,5; −1.51±1.18, 0.11±0.46, 0.02±0.32; p=0.007, n=6, n.s.; H: n=5,6,7; −1.51±1.18, −1.43±1.02, −1.94±1.35; p=0.71).

To examine hippocampal activity, a 32-channel linear array was inserted into CA1 of dorsal hippocampus (Fig. 1D). As previously described, neuronal firing was largely restricted to the pyramidal cell layer and occurred almost entirely during early sharp waves (eSPWs), a developmentally-transient burst driven by cortical input transmitted through the entorhinal cortex (Leinekugel et al., 2002; Mohns and Blumberg, 2010; Valeeva et al., 2019) (Fig.1E). Only animals with viral expression surrounding the recording electrode and limited to hippocampus were analyzed. Suppressing the excitability of hippocampal GABAergic neurons by subcutaneous injection of SalB decreased multi-unit activity (MUA) in the pyramidal cell layer relative to pre-injection baseline (Fig.1F). The amplitude and occurrence, but not the duration, of eSPWs, and the local field potential (LFP) spectra in a broad frequency range were also reduced (Fig.1F, Fig.S1C). By contrast, in different animals, enhancing GABAergic neuron excitability by injecting CNO increased pyramidal layer MUA but did not alter eSPW amplitude or LFP power (Fig.1G, Fig.S1A-B). Neither saline injection into animals expressing KORD/hM3Dq, nor SalB or CNO injections into GFP-only mice, altered activity. Thus, GABAergic interneurons excite the hippocampal CA1 network, and this excitation amplifies spontaneous activity during early development.

Such net-excitatory action of GABAergic neurons was no longer observed at P7 (Fig. 2). At P7, decreasing GABAergic neuron excitability increased pyramidal layer firing rates and the power of 6-14Hz frequencies in the LFP without significantly changing the occurrence, duration or amplitude of eSPWs (Fig. 2A-B, Fig.S2A). Enhancing interneuron excitability decreased pyramidal-layer firing and reduced LFP power in a broad frequency range (Fig.2C) without significantly affecting the eSPW (Fig.S2B). At P11, modulating GABAergic neuronal activity had similar effects on firing rates and LFP power (Fig.2D-E). These results demonstrate a reversal of hippocampal GABAergic interneuron function, from excitatory to inhibitory, between P3 and P7.

In contrast to hippocampus, reducing the excitability of cortical GABAergic interneurons at P3 increased MUA firing rates recorded within the injected area (Fig.3D-E), while increasing their excitability had the opposite effects (Fig.3G). Neither manipulation significantly changed LFP spectral power, nor the occurrence, duration, or amplitude of spontaneous spindle-bursts (Fig.S3), the cortical activity driven by spontaneous retinal activity at this age (Hanganu et al., 2006). As in CA1, neither saline injection in DREADD expressing animals, nor active drug in GFP expressing animals, had an effect. Modulation of GABA neuron excitability at P7 and at P11 had similar effects (Fig.3F, H). Thus, GABAergic interneurons have a net inhibitory role in visual cortex at ages at which they are excitatory in hippocampus.

To verify that the regional heterogeneity in DREADD response was a direct result of activity modulation of the interneurons, we used a mechanistically independent means to reduce GABAergic neuron firing. Photostimulation of JAWS, a light-driven inward chloride pump (Chuong et al., 2014), virally expressed in GABAergic interneurons hyperpolarized the membrane potential of GABAergic neurons in slices at all ages. Optogenetic suppression of GABAergic neuron activity *in vivo* at P3 and P7 showed similar effects to DREADD-based suppression (Fig. S4).

Such regional heterogeneity of interneuron function suggests that a systemic modulation of GABAergic activity, as used in epilepsy treatment, would produce diverse outcomes. To test this, we expressed KORD and hM3Dq in hippocampus as well as cortex, which provides the major drive to CA1 at these ages (Mohns and Blumberg, 2010; Valeeva et al., 2019), and recorded activity in the ipsilateral CA1 at P3. In contrast to injections limited to hippocampus, adding cortical suppression of GABAergic neuron excitability increased CA1 firing, while its enhancement reduced CA1 firing (Fig. S5). Thus at P3, inhibition of interneurons in cortex can overwhelm local excitation by interneurons in hippocampus, causing a net inhibitory effect in CA1.

The regional heterogeneity of GABAergic neuron excitation might be a direct result of differences in the excitation caused by the GABA_A_ receptor-mediated anion conductance, or by differential network effects such as desynchronization of local firing by depolarization (Spoljaric et al., 2017; Valeeva et al., 2010). To differentiate these possibilities we directly assayed the effect of activating anion conductance on pyramidal neurons in hippocampus and cortex using stGtACR2, a light-gated anion channel with a soma-targeting motif (Mahn et al., 2018), virally expressed in EMX1-cre mice (Gorski et al., 2002) (Fig. 4A, B). *In vivo* photostimulation of stGtACR2 in glutamatergic neurons of VC for 1 second reduced firing at both P3 and P7 (Fig. 4C), indicating an inhibitory role for anions in cortex at these ages as in adults (Mahn et al., 2018). In CA1 hippocampus at P3, as predicted by the manipulation of GABAergic neurons (Fig.1, S5), photostimulation of stGtACR2 in glutamatergic neurons induced immediate firing in the pyramidal cell layer (Fig. 4C). At P7, stGtACR2 activation became strongly inhibitory (Fig. 4C). Thus the heterogeneity of interneuron roles between regions likely reflects the differential intracellular concentration of anions, particularly chloride.

**Fig. 4.**
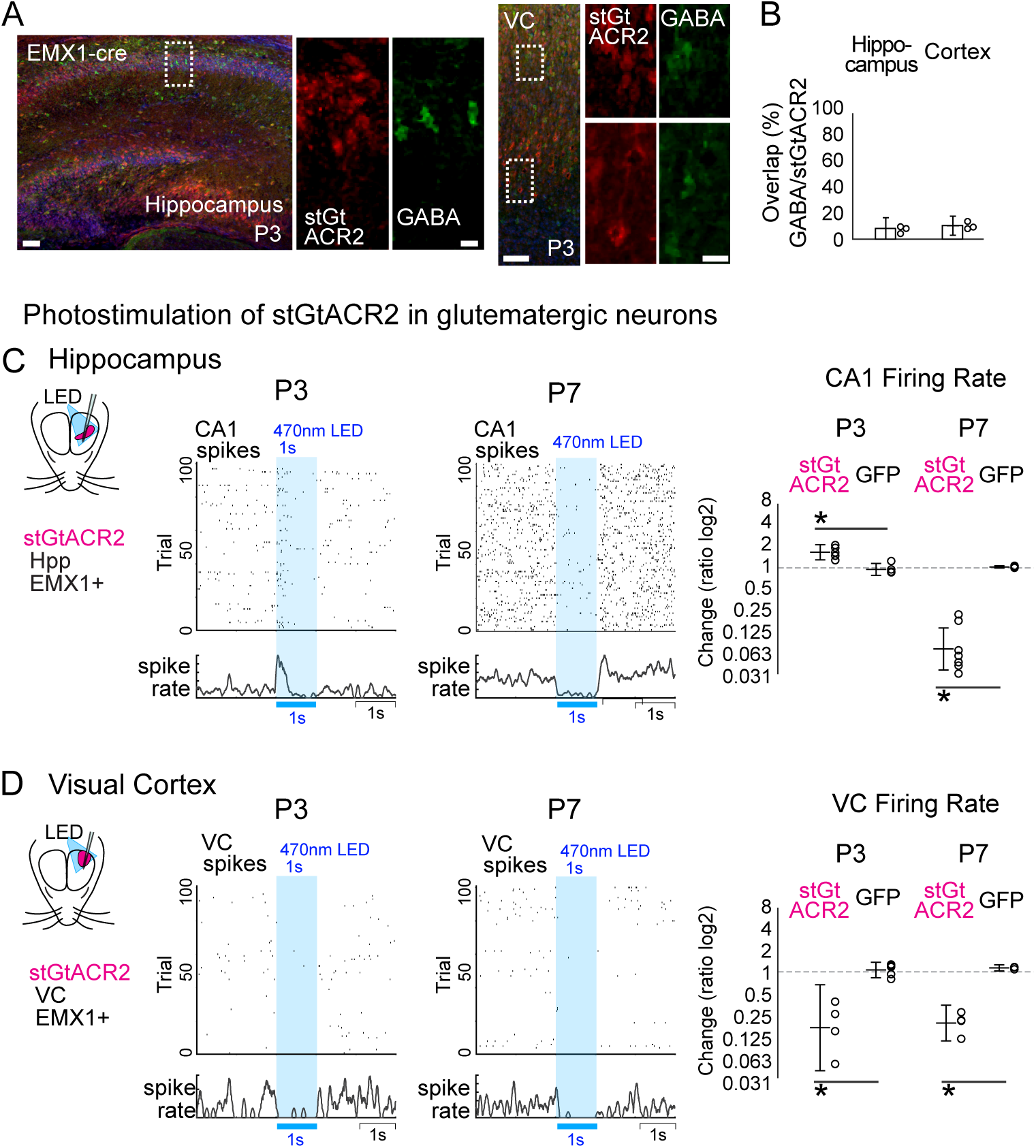
Changing role of anion conductance likely underlies the regional and age heterogeneity of GABA function. **(A, B)** stGtACR2, an anion-conducting channelrhodopsin with a soma targeting motif, is virally expressed in non-GABAergic neurons in hippocampus or cortex of EMX1-cre mice (n=3, 3; 8.5%±8.5%, 10.7%±7.6%). Scale bars: 50μm and 10μm. **(C)** Photostimulation of stGtACR2 in hippocampal glutamatergic neurons (470 nm LED, 1 sec) increased CA1 firing at P3 but decreases it at P7 (n=5,5,7,5; 0.74±0.38, −0.07±−0.35, −3.51±−4.25, 0.04±−0.01; p=0.0011, p<0.0001). **(D)** In visual cortex, photostimulation of stGtACR2 in glutamatergic neurons decreased MUA at both P3 and P7 (n=4,5,4,3; −2.65±−4.68, 0.08±−0.29, −2.43±−3.27, 0.18±0.04; p=0.0022, p=0.0004).

For this study, we have overcome a major technical barrier to manipulation of GABAergic neuron activity in awake neonatal mice as early as P3. This allowed us to investigate the role of interneurons in endogenous spontaneous activity fundamental to circuit formation. The results provide direct evidence that GABA can be excitatory *in vivo*, and that this excitatory drive is mediated by the activity of local interneurons which increase spiking in CA1. Under the same experimental conditions, these data also show that the net excitatory effect of local interneurons is more developmentally and regionally restricted to cortical circuits than would be predicted by *ex vivo* studies (Ikeda et al., 2003; Rivera et al., 1999). Such heterogeneity has been reported between cortical and sub-cortical regions of the same sensory system (Glykys et al., 2009; Ikeda et al., 2003) as well as within regions depending on neuronal birth-date (Yamada et al., 2004), and even within neurons between the soma and axons (Mahn et al., 2018), but cortical circuits have been widely assumed to share a similar developmental trajectory.

Our results show a strong dependence of excitability on GABAergic interneurons in developing hippocampus and cortex, but contributing minimally to network dynamics in the thalamocortical circuit. This is consistent with *ex vivo* studies showing a dependence of hippocampal spontaneous activity on GABA, while cortical activity requires glutamatergic transmission (Garaschuk et al., 2000). This may indicate a fundamental difference in the role for inhibitory circuits in each region and the computations each preform. In thalamocortical circuits, excitatory drive and early oscillations are generated by the relay thalamus and its loop with cortex (Colonnese and Phillips, 2018), and inhibition serves to restrict activity topographically (Minlebeav et al 2007). By contrast topography could be less relevant in the hippocampus, where early activity is coordinated by GABAergic hub cells (Bonifazi et al 2009).

By showing that the textbook model of excitatory GABA does occur *in vivo*, and that local interneurons are critical to the generation of early hippocampal activity, though in a much more heterogeneous group of regions and ages than previously expected, our results support the likely hypothesis that regulation of the chloride reversal potential by hormones (Tyzio et al., 2006), epilepsy (Marguet et al., 2015) and autism risk (He et al., 2014; Tyzio et al., 2014) may have important effects on developing networks; though our results also show it is essential to test these effects *in vivo*.

## Materials and Methods

### Experimental design

We employed chemogenetic and optogenetic approaches combined with cre-line mice to specifically manipulate activity of GABAergic neurons (GAD2-cre mice: suppression by KORD with its ligand SalB, enhancement by hM3Dq with its ligand CNO; suppression by JAWS with green-yellow LED) and also to activate anion channels in glutamatergic neurons (EMX1-Cre mice: stGtACR2 with blue LED). The effects of chemogenetic and optogenetic manipulation on endogenous network activity were monitored using multi-electrode array recordings in un-anesthetized neonatal mice.

### *In vivo* recording

All experiments were conducted with approval from The George Washington University School of Medicine and Health Sciences Institutional Animal Care and Use Committee, in accordance with the *Guide for the Care and Use of Laboratory Animals* (NIH).

GAD2-IRES-Cre (Gad2^tm2(cre)Zjh^, Stock No: 010802) (Taniguchi et al., 2011) and EMX1-IRES-Cre (B6.129S2-Emx1^tm1(cre)Krj/J^, Stock No: 005628) (Gorski et al., 2002) mice were acquired from The Jackson Laboratory. C57BL/6 mice were acquired from The Jackson Laboratory and Hilltop Lab Animals. All GAD2-cre and EMX1-cre mice used were heterozygous, obtained by crossing homozygous GAD2-cre or EMX1-cre males and C57BL/6 females. Animals were housed one litter per cage on a 12/12 light/dark cycle. Both males and females were used. *In vivo* recording methods are described previously (Colonnese et al., 2017; Murata and Colonnese, 2016, 2018). Topical lidocaine (2.5%) and systemic Carprofen (5 mg/kg) were used for preoperative analgesia. To place the headplate, under isoflurane anesthesia (3% induction, 0.5–1.5% maintenance, verified by toe pinch), the scalp was resected, the skull cleaned, and a stainless plate with a hole was placed so that the region over occipital cortex and hippocampus was accessible. The plate was fixed to the skull with vetbond and dental cement. Pups were monitored for signs of stress after recovery from anesthesia. For recording, the animal was head-fixed, and the body was supported within a padded enclosure. Body temperature was monitored with a thermocouple placed under the abdomen and maintained at 34–36°C by heating pad placed under the body restraint. Body movement was detected using a piezo-based detector placed under the enclosure. Electromyogram (EMG) was recorded from the ventral neck by a single stainless steel wire electrode. An Ag/AgCl wire was placed through the skull over cerebellum as ground.

A variety of single and multi-shank silicon polytrodes (Neuronexus) were used, including 32-channel linear “edge” arrays with 100, 50, or 20 μm contact separation, and the “Poly2” configuration with two parallel rows with 50 μm separation. Electrodes were coated with DiI (Life Technologies) before insertion for histological verification of electrode location. For VC recording, the monocular primary visual cortex was targeted with the following coordinates: −0.2~+0.2 mm anterior, 1.5~2.1mm lateral from the lambda at P3; −0.2~+0.2mm and 1.7~2.3mm at P7; and −0.2~+0.2mm and 2.1~2.9mm at P11. For hippocampal recording, the dorsal CA1 was targeted with the following coordinates: 0.5~0.8 mm anterior and 1.3~1.7mm lateral from the lambda at P3; 0.7~1.1mm and 1.6~2.0mm at P7; and 0.9~1.5mm and 1.7~2.3mm at P11.

SalB (1mg/kg, 0.1mg/ml in saline with 1% dimethyl sulfoxide (DMSO)) was subcutaneously injected to activate KORD, and CNO (10mg/kg; 1mg/ml in saline with 1% DMSO) was subcutaneously injected to activate hM3Dq (Vardy et al., 2015). Saline with 1% DMSO was subcutaneously injected as control.

For activating JAWS, a 400 μm diameter optic fiber (Thorlabs, M128L01) coupled with green-yellow LED (Thorlabs, MINTF4, peak at 554nm, 80% intensity at 520nm-586nm, 21.2mW at the fiber tip) was placed above the skull over the dorsal hippocampus or visual cortex. 1 second LED stimulation was given at every 15-20 seconds. For activating stGtARC2, photo-stimulation using 470nm LED (Thorlabs, M470F3, peak at 467nm, 80% intensity at 464nm-471nm, 12.5mW at the fiber tip) were similarly performed.

Electrical signals were digitized using the Neuralynx Digital Lynx S hardware with Cheetah v5.6 software. dEEG signals were bandpass filtered between 0.1 Hz and 9 kHz, and digitized at 32 kHz. Cortical recordings were referenced to a contact site located in the underlying white matter. Hippocampus recordings were referenced to a contact just dorsal to hippocampus. Multiunit activity (MUA) was extracted by threshold crossing of −40 (P3) or −50 (P7 & P11) μV following 300 Hz to 9 kHz bandpass filtering.

### Analysis

Neural signals were imported into MATLAB (MathWorks). Spike-times and dEEG were downsampled to 1 kHz. Before analysis, six animals were excluded for unstable baseline LFPs (periods with LFP amplitudes larger than the maximum amplitude of visual response or with electrical or movement artifacts) or spike activity (more than 20% changes between the start and end of recording), eight animals were excluded for viral injection spread outside the recording region (hippocampus or visual cortex), and five animals were excluded because post-recording histology showed the electrode was not located in CA1 of the hippocampus.

For hippocampus, the pyramidal cell layer was identified in each recording as the channel with the highest MUA spike rate, and the *strata radiatum* was identified as the channel with largest negative LFP deflection. Average distance of the *strata radiatum* from the pyramidal cell layer was ~300μm at all ages. eSPWs were detected by identifying a triggering threshold of root mean square (RMS, 9ms window) of the strata radiatum LFP at 7 standard deviations (std) above the mean. The beginning and end of the eSPW is defined as the points when the RMS remained above 2 std and contained the initiation threshold.

For cortex, at P3 & P7, presumptive layer 4 was identified in each recording as the layer with the largest negative LFP deflection during spindle-bursts (Colonnese and Khazipov, 2010). In P11 animals, cortical layer 4 was identified in each recording as the channel with the earliest negative LFP deflection and the fastest spike response in the mean visual evoked response as previously described. (Colonnese and Khazipov, 2010). Spindle-bursts were identified by at least one negative trough of the layer 4 LFP that was greater than 5 std from the mean; additional cycles with negative LFP greater than 2 std within 1 second of each other were used to define the length of the burst.

For all analyses, after recovery period for at least 30 min following electrode insertion, baseline activity was calculated from the entirety of a continuous 20 min period, while the experimental condition (KORD-SALB, KORD-saline, GFP-KORD, hM3Dq-CNO, hM3Dq-saline, GFP-CNO) was calculated from a 20 min period beginning 10 min after injection. For spectral analysis, LFP spectra from presumptive layer 4 (visual cortex) or *strata radiatum* (hippocampus) were obtained by averaging 2s multi-taper windows (time-band width 3 with 5 tapers (Chronux package (Mitra and Bokil, 2007)). To reduce the effect of the 1/*f* relationship, mean multitaper spectra were multiplied by frequency. The frequency axis was resampled on a log scale to equalize the representation of high and low frequencies and reduce the multiple-comparisons problem. For normalization, frequency power at each band was divided by the mean 1-100 Hz power of baseline. For optogenetic stimulation, spike rates for baseline (900ms duration prior to LED stimuli) and for during photostimulation (900ms duration starting 20ms after the onset of LED stimuli to exclude potential photo-artifacts and antidromic spikes by stGtACR2 (Mahn et al., 2018)) were analyzed.

### Virus injection

AAV8-hSyn-dF-HA-KORD-IRES-mCitrine (65417-AAV8, tier >7e12 GC/ml) (Vardy et al., 2015), AAV8-hSyn-DIO-hM3D(Gq)-mCherry (44362-AAV8, tier >1e13 GC/ml), AAV8-hSyn-DIO-hM3D(Gq)-IRES-mCitrine (50454-AAV8, tier >1e13 GC/ml))(Armbruster et al., 2007; Krashes et al., 2011), and AAV1-hSyn1-SIO-stGtACR2-FusionRed (105677-AAV1, tier >1e13 GC/ml) (Mahn et al., 2018) were obtained from Addgene, and AAV1-CAG-FLEX-GFP(AAV1-AllenInstitute854, tier >2.9e13 GC/ml) and AAV1-CAG-FLEX-tdTomato (AAV1-AllenInstitute864, titer 8.52e11 GC/ml) were obtained from The Penn Vector Core. AAV8-CAG-FLEX-rc-[Jaws-KGC-GFP-ER2] (Addgene, plasmid# 84445) (titer 8.5e12 GC/ml) (Chuong et al., 2014) was obtained from the UNC Vector Core (Courtesy of Dr. Chuong and Dr. Boyden).

Mouse pups at the day of birth (P0) were cold anesthetized, and 30–100 nl of viral solution was injected locally into the hippocampus (0.4-0.8mm anterior and 1.2-1.8mm lateral from the Lambda, 1.0-1.5mm depth) or visual cortex (−0.2-0.0 mm anterior and 1.5-2.0mm lateral from the Lambda, 0.4-0.7mm depth) using a Nanoject II (Drummond) (Murata and Colonnese, 2016). Two days post injection yielded visible expression of fluorophore around the injected sites.

Expression local to the recording electrode and limited to the region of interest was verified in all animals recorded. For chemogenetic manipulation, about the half of animals were injected with only AAV-FLEX-GFP for control, and the rest were injected with AAV-dF-KORD and/or AAV-DIO-hM3Dq and/or AAV-FLEX-GFP. Similarly, for optogenetic manipulation, about the half animals were injected only with AAV-FLEX-GFP or AAV-FLEX-tdT and the rest were injected with AAV encoding opsins. Recordings and analyses were conducted in a blind manner.

### Slice recording

Slice recordings were conducted as previously described with slight modifications (Bolton et al., 2015; Zhao et al., 2013). Mice were anesthetized with Isoflurane and decapitated. The brain was quickly removed and placed in ice-cold cutting solution containing 110 mM Choline Chloride, 2.5 mM KCl, 25 mM NaHCO_3_, 1.25 mM NaH_2_PO_4_, 12 mM glucose, 3 mM sodium pyruvate, 10 mM sodium ascorbate, 7 mM MgSO_4_, and 0.5 mM CaCl_2_. Coronal cortical and hippocampal slices (300 μm) were cut using a Leica Vibratome VT 1000S. The slices were kept in ACSF containing 127 mM NaCl, 2.5 mM KCl, 25 mM NaHCO_3_, 1.25 mM NaH_2_PO_4_, 12 mM glucose, 1 mM MgCl_2_, 2 mM CaCl_2_ at 32 °C for at least 15 min then at room temperature (22 - 25 °C). Slices were placed in a recording chamber at room temperature and continuously perfused with carbogenated ACSF. pClamp (Molecular Devices) was used to acquire and analyze data. Electrical signals were amplified with an Axopatch 200B amplifier, digitized with a Digidata 1322A interface (Molecular Devices), filtered at 2 kHz, sampled at 5 kHz.

Glass pipettes (2 - 6 MΩ) were filled with an internal solution containing the following: 105 mM K-gluconate, 10 mM Na-phosphocreatine, 4 mM EGTA, 10 mM HEPES, 4 mM Na-ATP, 1 mM Na-GTP, 3 mM MgCl_2_, 0.07 mM CaCl_2_, brought to ∼290 mOsm with ∼28 mM sucrose and pH 7.3 with KOH. GABAergic cortical and hippocampal neurons with fluorescence were patched under a 40× objective, and whole-cell current-cramp recordings were conducted to measure the changes in resting membrane potential following bath application of 100nM SalB or 100nM CNO. At least 1 minute of stable membrane potential was analyzed using Clampfit software (Molecular Devices) before and after treatment. For activating JAWS, 1 second photostimulation with green-yellow LED (Thorlabs, MINTL5, peak at 554nm, 14.2mW) was given for every 5 seconds for 10 times and similarly analyzed.

### Immunohistochemistry

Animals were perfused and post-fixed with 4% Paraformaldehyde in PBS. Brains were sectioned by vibratome at 150μm in the coronal plane. For cryosectioning, brains were cryoprotected in 10% and 30% sucrose in PBS, then sectioned at 40μm. The following antibodies were used: rabbit anti-GABA (Sigma, A2052, Lot# 103M4793), chicken anti-GFP (Abcam, ab13970, Lot# GR3190550-12), chicken anti-RFP (Rockland, 600-901-379S, Lot 42717), anti-chicken Alexa 488, anti-chicken Alexa 555, anti-rabbit Alexa 488, anti-rabbit Alexa 555, and DAPI (ThermoFisher). Confocal images were taken with Zeiss710 using a 10× objective, and analyzed with FIJI (ImageJ).

### Statistical Analysis

Mean ± 95 % confidence interval are reported, and individual data were presented except for spectra. Hypothesis tests were conducted using nonparametric methods when *n* < 10. One-way ANOVA was used for all tests of manipulation and age dependence, and *post hoc* test (Tukey HSD) used to examine differences between specific manipulation and age groups. Significant differences by *post hoc* test (*p* < 0.05) are reported as asterisks on the relevant figure. *P* values <0.001 are rounded to nearest power of 10. Spectra were examined at each frequency for significant difference using nonparametric permutation tests corrected for multiple comparisons by the method of Cohen (Cohen, 2014) with *p* < 0.05. All tests were performed in MATLAB. The number of animals, the statistical test, results and p-values are reported in the supplemental table.

## Acknowledgments

We thank Dr. David Mendelowitz for the slice electrophysiology equipment and Dr. Amy Chuong and Dr. Ed Boyden for AAV-FLEX-JAWS. We thank Dr. Chris McBain and lab, Dr. Leo Chalupa and Dr. Yehezekel Ben-Ari for comments on the project and manuscript. We thank Dr. Marnie Phillips for editing and composition on the manuscript.

## Funding

This work was supported by NIH Grant R01 EY022730 (M.T.C.).

## Author contributions

Y.M. designed the study and conducted the experiments. Y.M. and M.T.C. analyzed the results and wrote the manuscript.

## Competing interests

The authors declare no competing interests.

## Data and materials availability

The datasets generated and analyzed during the current study are available from the corresponding authors upon reasonable request.

---

**Fig. S1.**
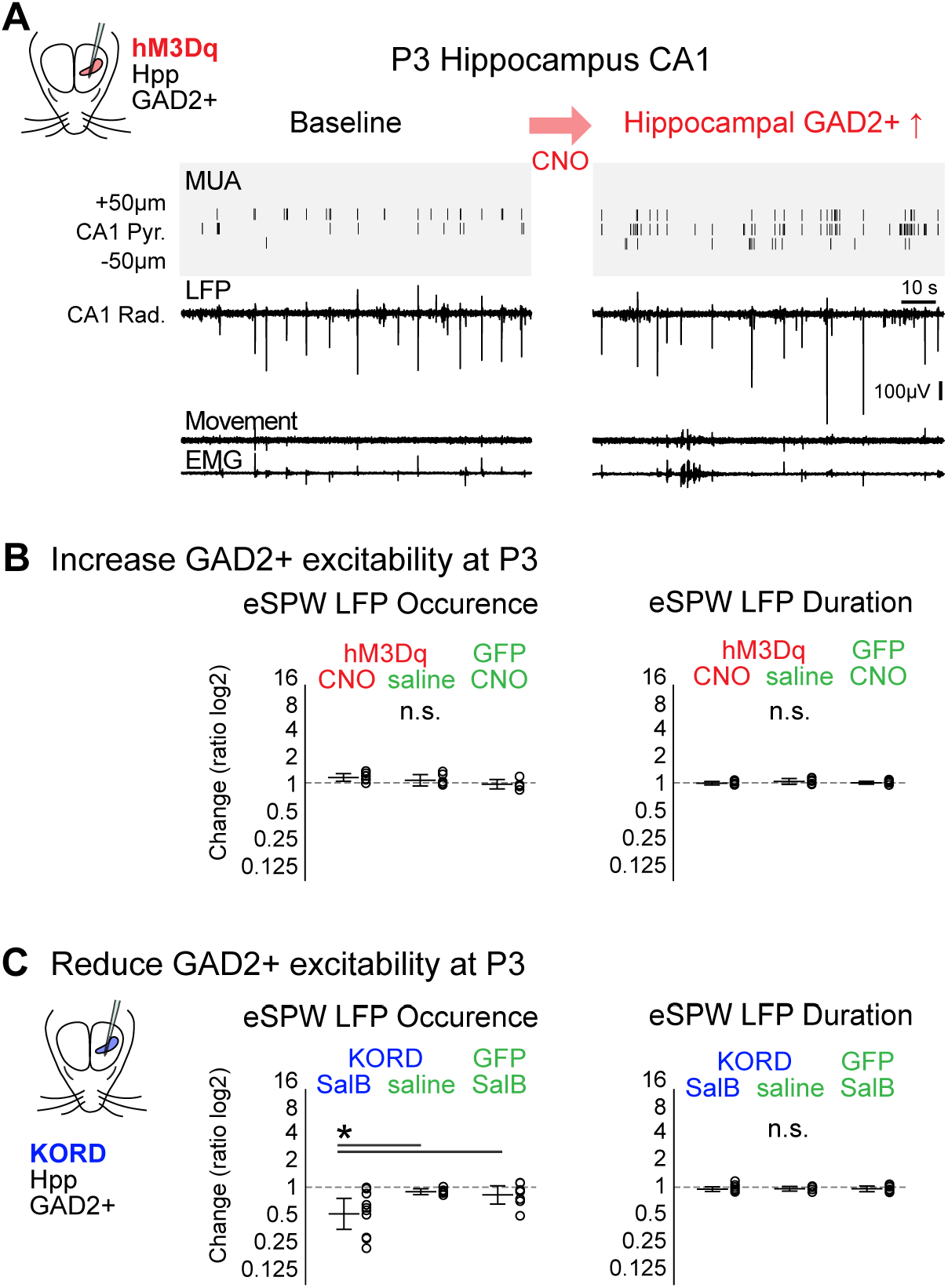
GABAergic neurons are excitatory in P3 hippocampus *in vivo*. **(A)** Representative recording of hippocampus at P3 and the effect of GABAergic neuron enhancement. **(B)** Enhancement of hippocampal GABAergic neurons increases MUA firing rates (Fig. 1G) without significantly changing the amplitude (Fig. 1G), occurrence, and duration of eSPWs. (n=7,6,6, 0.22±0.16, 0.11±0.23, −0.05±0.19, p=0.0631; −0.01±0.07, 0.06±0.12, 0.01±0.06, p=0.2937) **(C)** Suppression of hippocampal GABAergic neurons increase MUA (Fig. 1F), the amplitude (Fig. 1F) and occurrence but not duration of eSPWs. (n=10,6,8, −1.09±0.64, −0.18±0.12, −0.31±0.37, p=0.0166; - 0.08±0.1, −0.06±0.1, −0.07±0.11, p=0.9628)

**Fig. S2.**
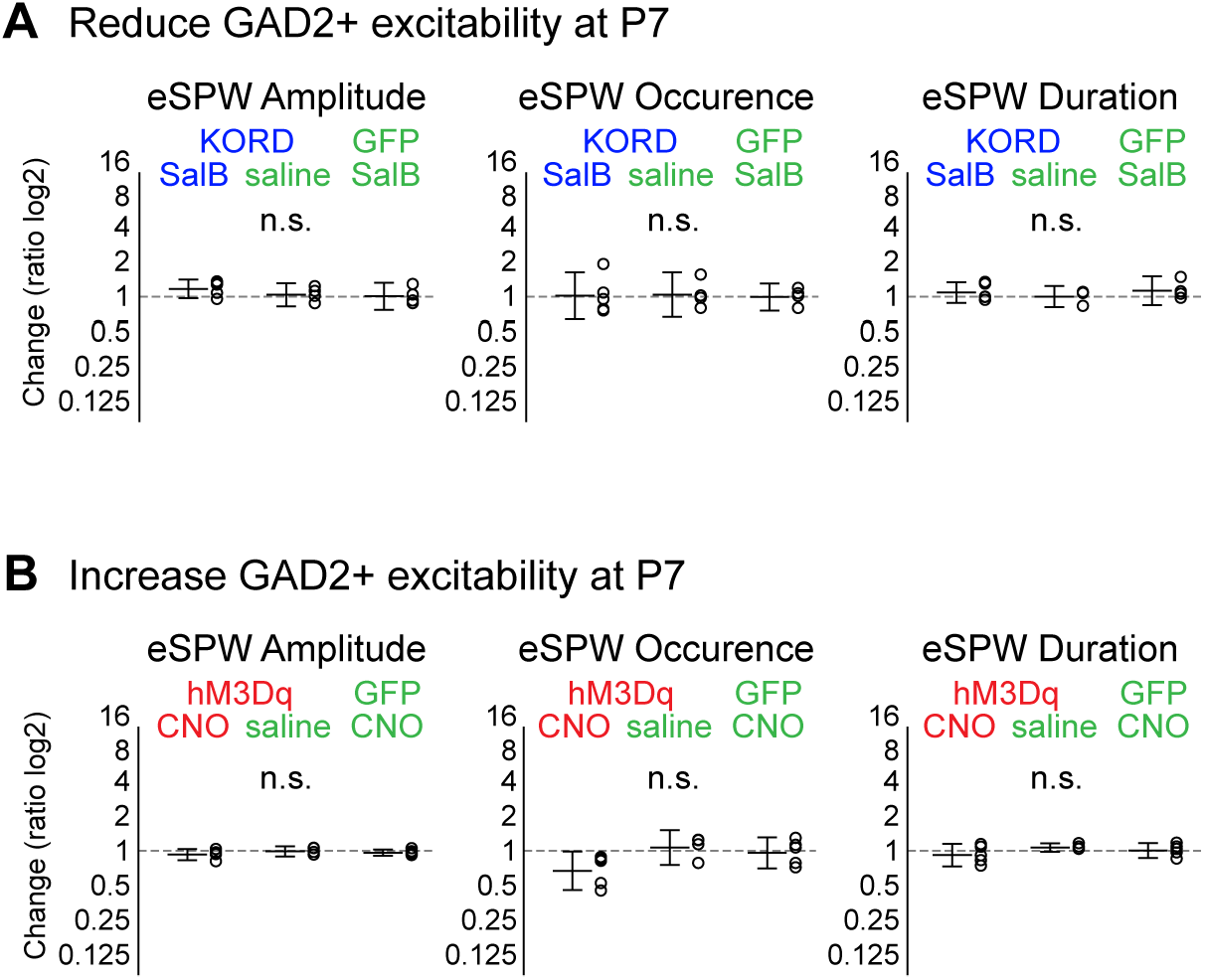
Hippocampal GABAergic neurons are inhibitory by P7. **(A)** Suppression of hippocampal GABAergic neurons at P7 increases MUA firing rates (Fig. 2B) without significantly changing the amplitude, occurrence, and duration of eSPWs. (n=5,4,4, 0.25±0.3, 0.06±0.37, 0.01±0.44, p=0.3491; 0.03±0.76, 0.06±0.72, −0.01±0.44, p=0.9754; 0.13±0.34, 0±0.34, 0.19±0.47, p=0.5855) **(B)** Enhancement of hippocampal GABAergic neurons at P7 increase MUA (Fig. 2C) without significantly changing the amplitude, occurrence and duration of eSPWs. (n=5,4,5; −0.12±0.18, −0.02±0.17, −0.06±0.1, p=0.4125; −0.65±0.63, 0.11±0.56, −0.06±0.5, p=0.0517; −0.14±0.36, 0.11±0.14, 0.01±0.24, p=0.2714)

**Fig. S3.**
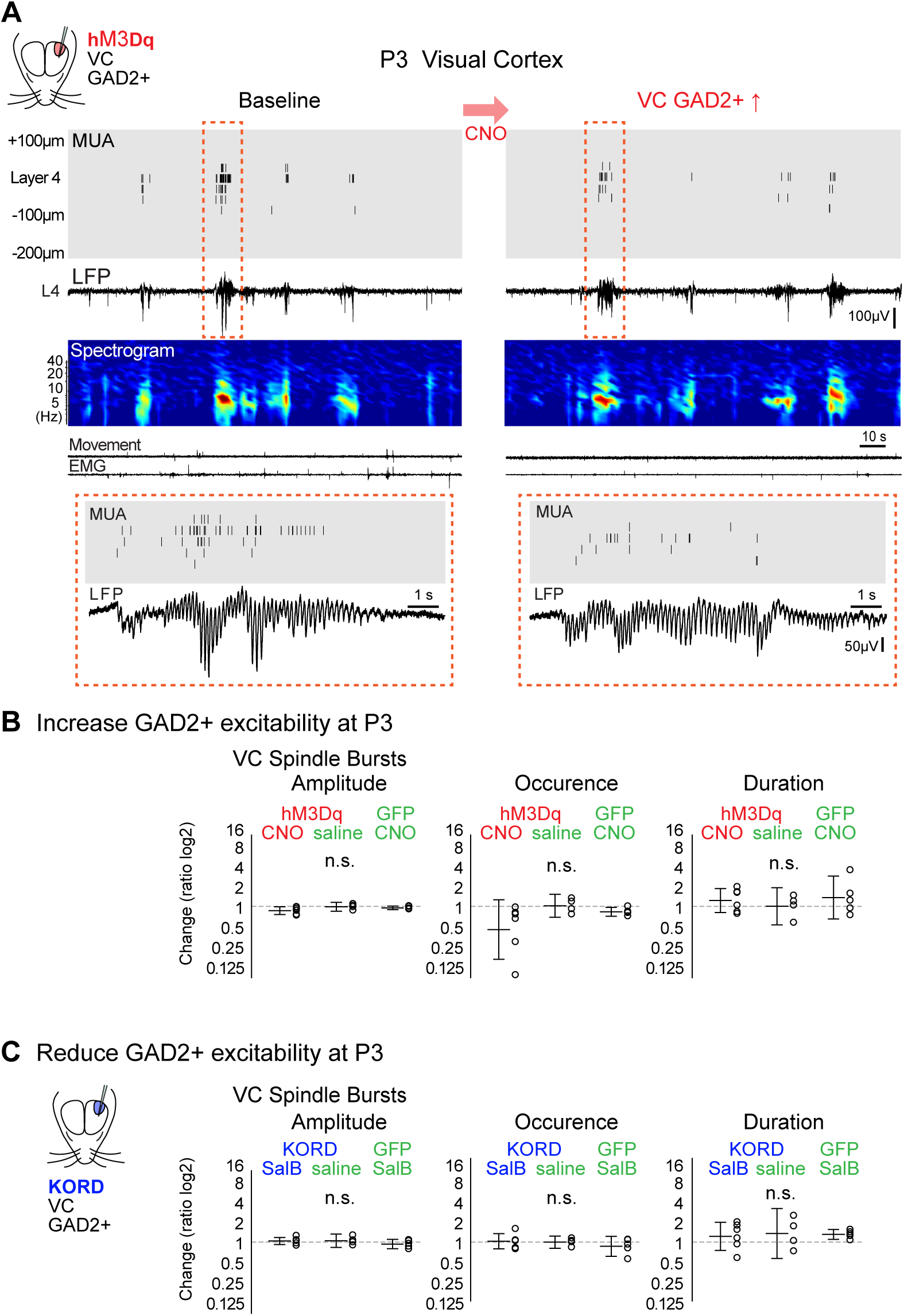
GABAergic neurons in visual cortex have a net inhibitory action at P3. **(A)** Representative recording of visual cortex at P3 and the effect of GABAergic neuron enhancement. **(B)** Enhancement of visual cortical GABAergic neurons decreases MUA firing rates (Fig. 3G) without significantly changing the amplitude, occurrence, and duration of spindle-burst oscillations. (n=6,4,5; −0.24±0.22, −0.02±0.25, −0.09±0.1, p=0.1267; − 1.3±1.67 0.03±0.65, −0.31±0.25, p=0.1495; 0.33±0.67, 0.01±1.04, 0.49±1.21, p=0.6455) **(C)** Suppression of visual cortical GABAergic neurons increase MUA (Fig. 3E) without significantly changing the properties of spindle-burst. (n=6,4,5; 0.06±0.19, 0.07±0.37, −0.11±0.26, p=0.3256; 0.04±0.43, −0.01±0.34, −0.24±0.56, p=0.4675; 0.32±0.79, 0.48±1.4, 0.43±0.28, p=0.9292)

**Fig. S4.**
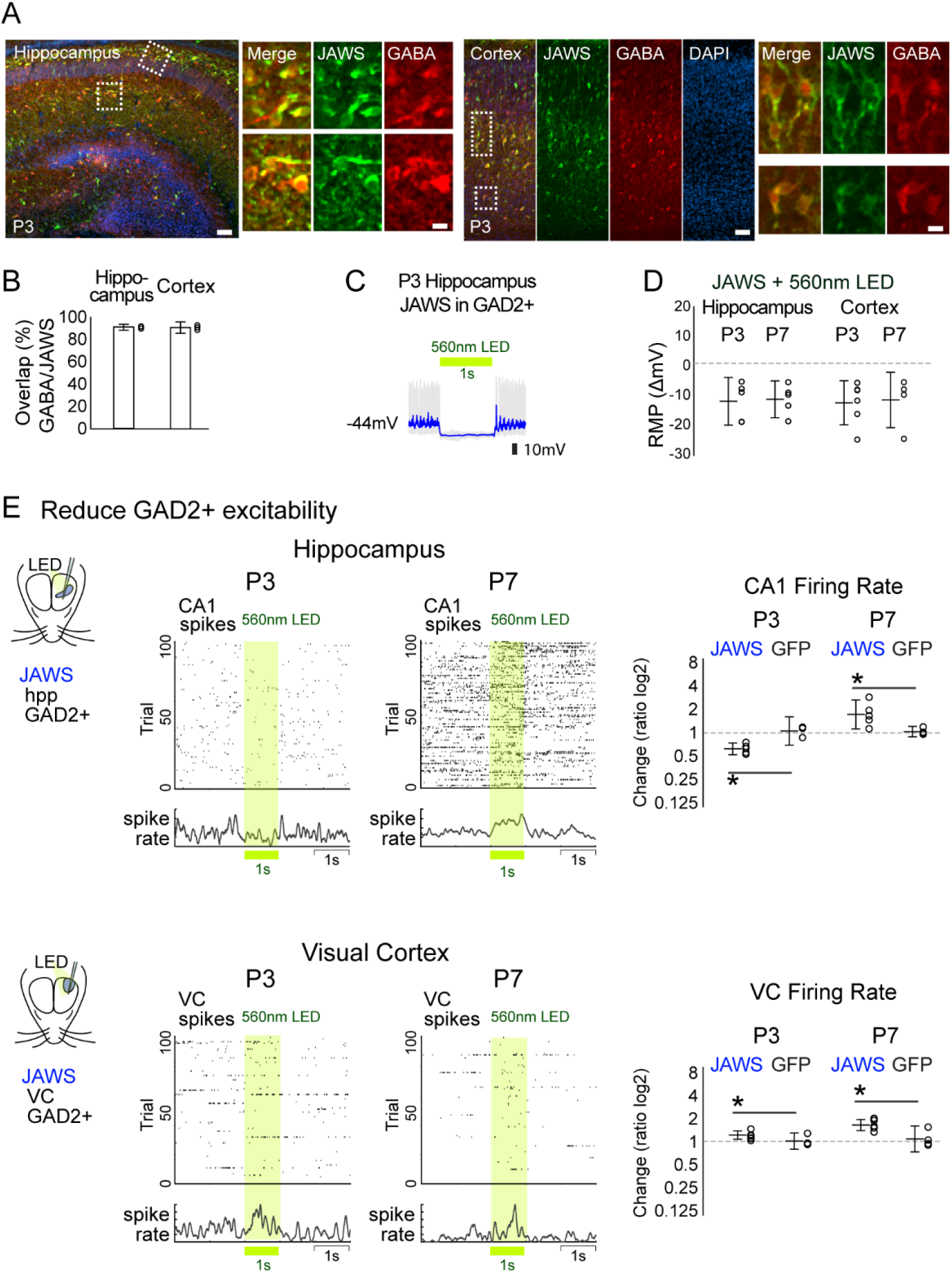
*In vivo* optogenetic suppression of GABAergic neurons demonstrates the excitatory-to-inhibitory shift of GABAergic neuron function between P3 and P7 in hippocampus but constantly inhibitory in cortex. **(A, B)** JAWS, light-gated chloride inward pump, is virally expressed in GABAergic neurons in hippocampus or cortex of GAD2-cre mice (n=3, 3; 91.2%±2.7%, 91.0%±5.1%). Scale bars: 50μm and 10 μm. **(C, D)** Changes in resting membrane potential caused by photostimulating JAWS-expressing GABAergic neurons in hippocampal and cortical slices at P3 and P7. (n=5,5,6,5; −11.82±8.02, −11.15±6.2, −12.33±7.43, −11.37±9.33). **(E)** (Top) *In vivo* optogenetic suppression of hippocampal GABAergic neurons, by photostimulating JAWS with green-yellow LED (560nm, 1sec), reduced CA1 MUA firings at P3 but instead increased it at P7 (n=5,4,5,7; −0.65±−0.89, 0.13±−0.05, 0.75±0.16, 0.05±−0.07; p=0.0002, p=0.0037). (Bottom) By contrast, in the visual cortex, optogenetic suppression of GABAergic neurons increased MUA firings at both P3 and P7 (n=6,6,7,4; 0.32±0.16, 0.01±−0.28, 0.66±0.44, 0.11±−0.44; p=0.0337, p=0.0117).

**Fig. S5.**
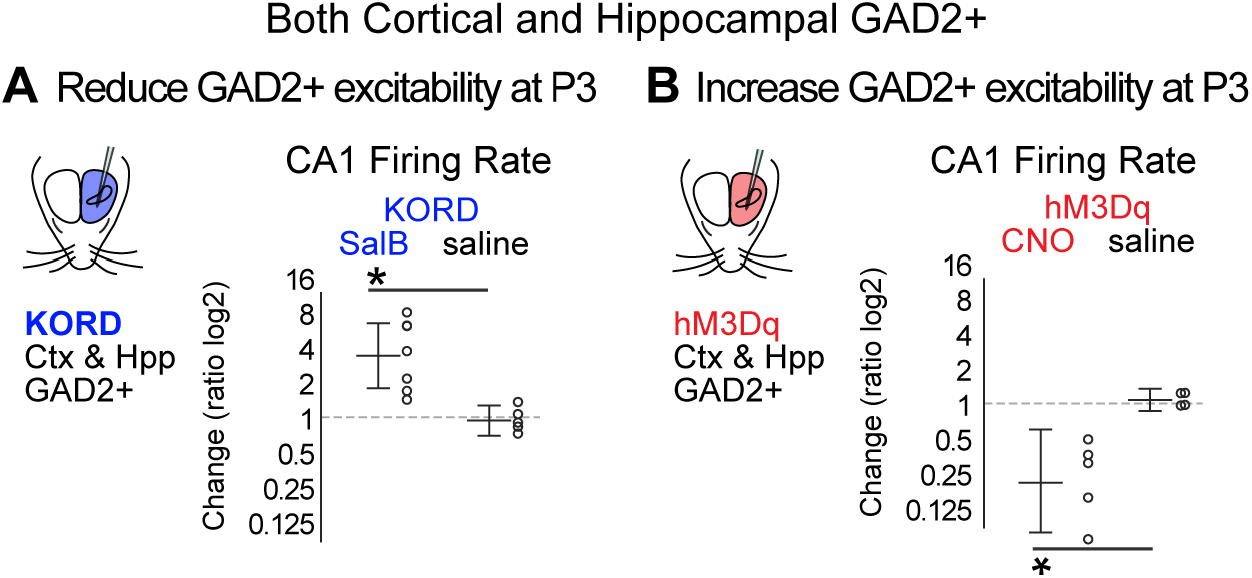
GABAergic inhibition in cortex swamps local GABAergic excitation in hippocampus. **(A)** Change in CA1 pyramidal cell firing following suppression of GAD2+ neurons expressing KORD in both hippocampus and cortex at P3 (n=7, 5; 1.98±0.94, −0.11±−0.59; p=0.003). **(B)** Activation by hM3Dq (n=5, 4; −2.56±1.85, 0.11±0.36; p=0.01). Note both changes are opposite that observed for local viral expression (Fig.1)

**Table S1.**
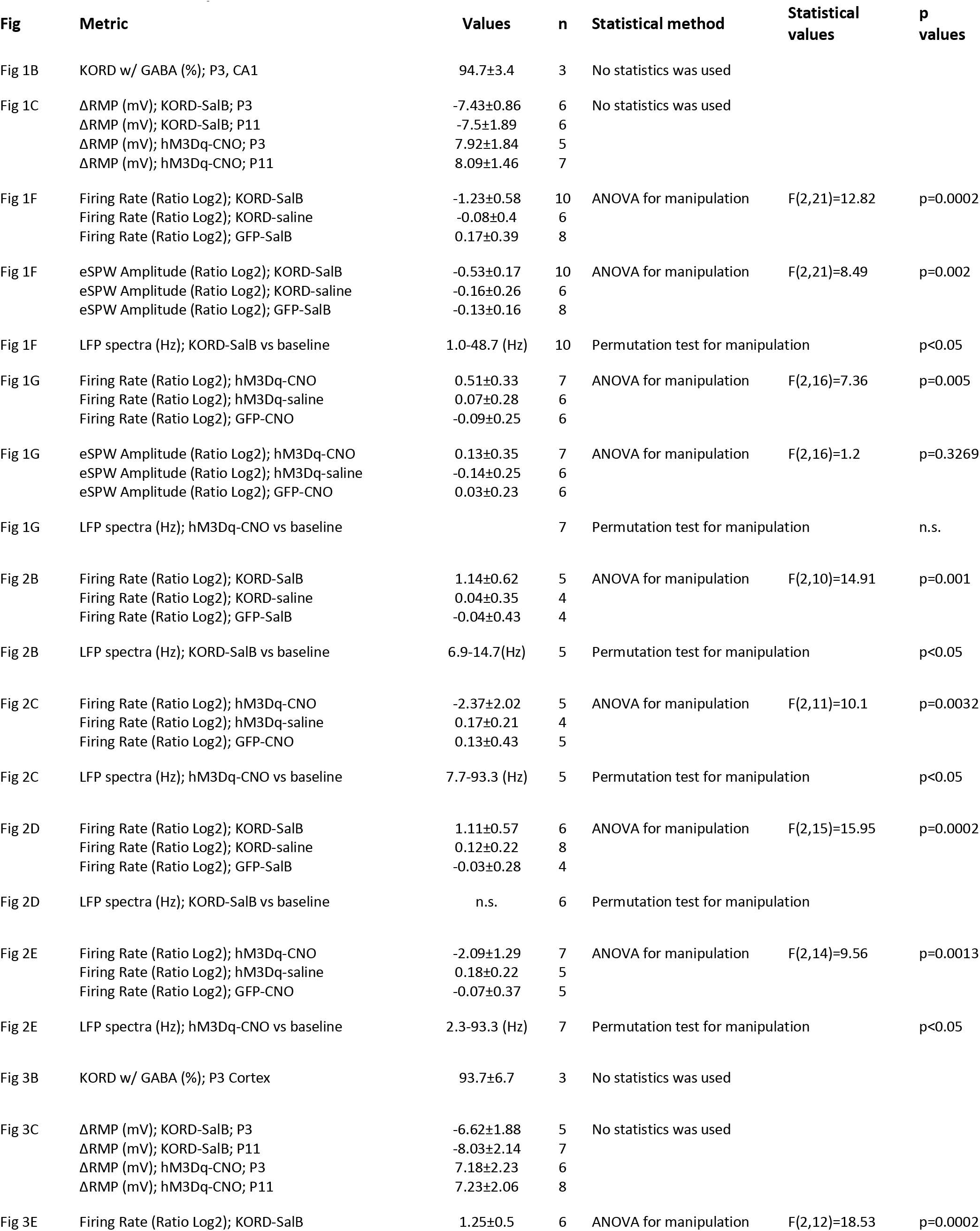
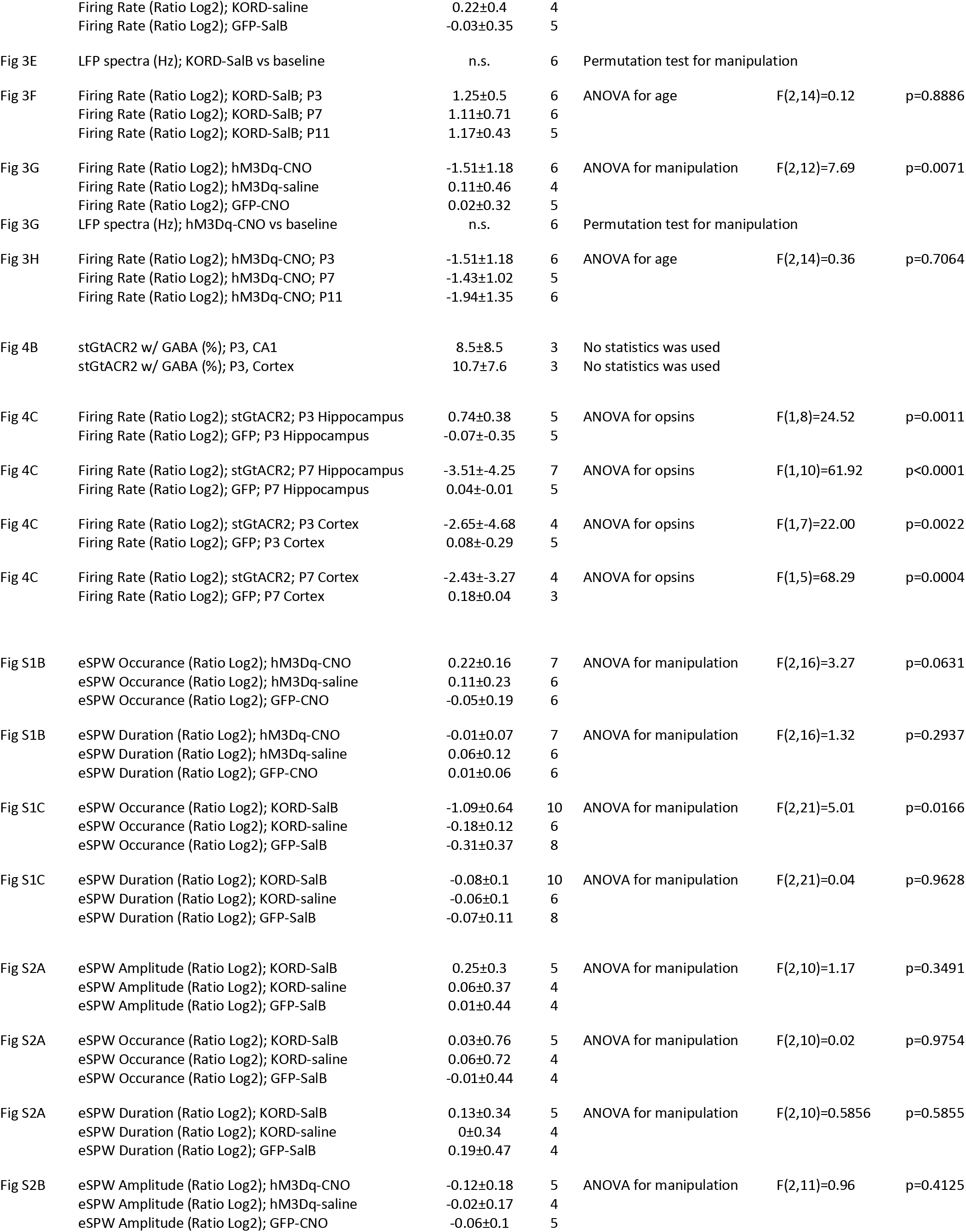
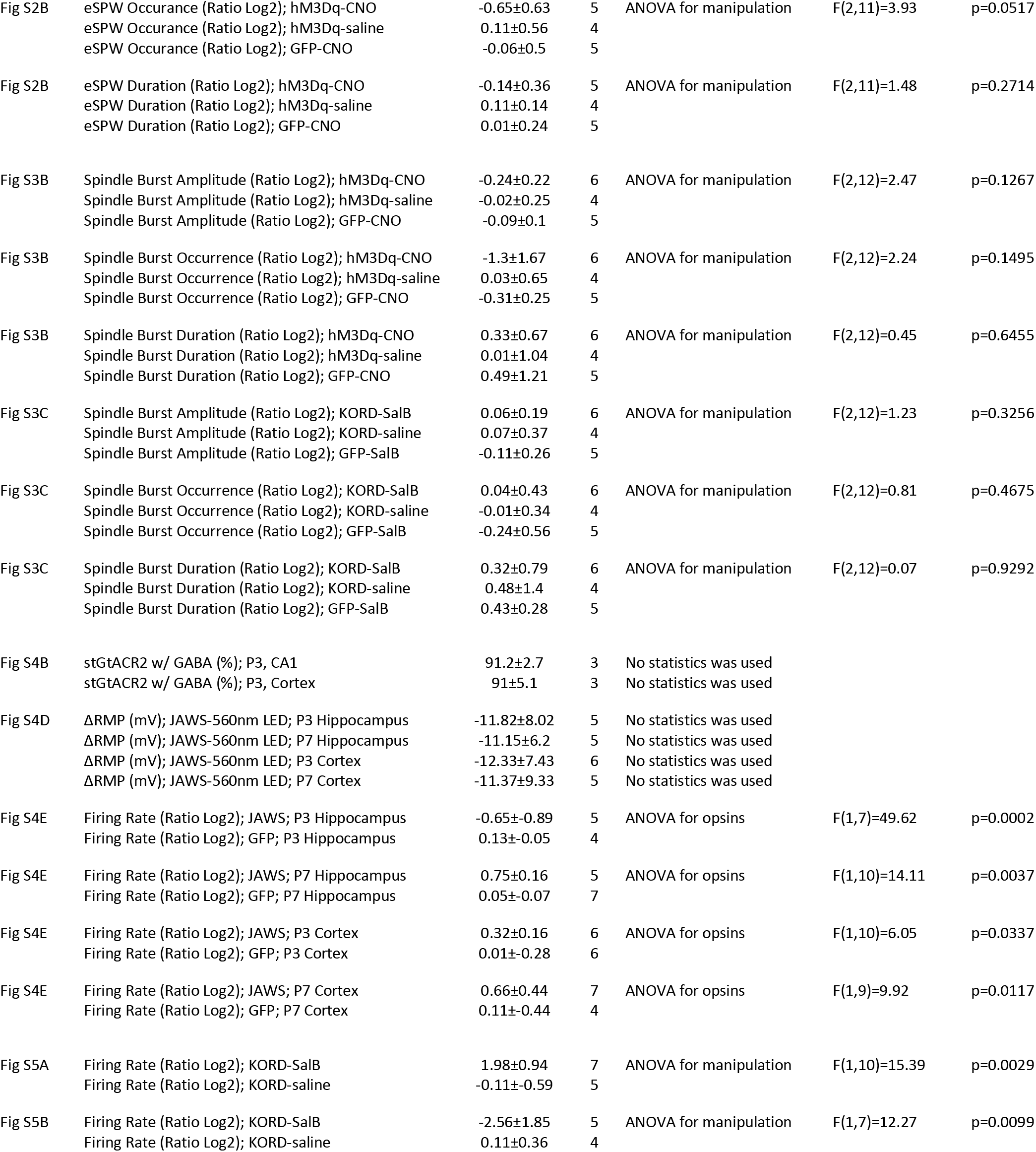
Summary data

